# Disentangling unique site-specific and shared habitat-level adaptation in a classic system of repeated evolution

**DOI:** 10.64898/2026.04.04.716503

**Authors:** Marius Roesti, Hannes Roesti, Hiranya Sudasinghe, Nicole Nesvadba, Verena Saladin, Catherine L. Peichel

## Abstract

Repeated divergence across contrasting habitats is widely used to infer natural selection and adaptation. However, such inferences remain inherently correlative and capture only adaptation shared among populations from the same habitat type, thereby missing site-specific adaptation unique to individual populations. Field transplant experiments test adaptation more directly by measuring fitness in nature, but they are typically limited to pairwise reciprocal exchanges between populations and therefore cannot separate shared habitat-level and unique site-specific adaptation. Here we extend the typical transplant framework to include multiple populations from both within and across habitat types, allowing fitness variation to be partitioned into shared habitat-level and unique site-specific components. We apply this framework to lake–stream stickleback, a classic system for studying adaptation via repeated divergence. Specifically, we transplanted laboratory-reared fish from a panmictic lake population and four independent stream populations across one lake and two stream sites. Stream fish outperformed lake fish in streams and vice versa, demonstrating adaptive lake–stream divergence. However, at both stream sites, local stream fish also outperformed foreign stream fish. Strikingly, this site-specific advantage was twice as large as the advantage of foreign stream fish over lake fish, reflecting the fitness benefit of shared stream adaptation. These results show that most fitness-relevant evolutionary variation in this system is unique to individual populations and therefore invisible to approaches that rely on repeated evolution to infer adaptation. More broadly, our work underscores the importance of ecological scale for understanding adaptation and evolutionary predictability.

**Significance statement:** Repeated evolutionary responses to similar environments commonly serve as evidence for adaptation. However, such comparative approaches do not directly measure fitness or resolve whether variation among populations from the same nominal habitat reflects adaptation to site-specific conditions. Using laboratory-raised lake and stream stickleback transplanted both within and across habitat types, we separated the fitness effects of shared adaptation to the same nominal habitat and site-specific adaptation unique to individual populations. Site-specific adaptation explained two-thirds of the total fitness variation, exceeding the contribution of adaptation shared among populations from the same habitat type. Our study highlights that natural selection generates fitness-relevant divergence at finer ecological scales than comparative approaches can resolve, underscoring the irreplaceable value of field experiments for understanding evolution.

## Introduction

Repeated evolutionary responses to similar ecological conditions are a hallmark of natural selection because non-adaptive processes are unlikely to generate consistent evolutionary outcomes (1–4). Accordingly, evolutionary biologists often test for adaptation by asking whether replicate populations in different habitats show repeated patterns of divergence. In such comparisons, the repeated component of divergence between habitats is treated as evidence for adaptation. However, this framework can only detect components of adaptation that differ consistently between habitats and are therefore shared among populations within the same habitat type. Yet even in classic systems for studying adaptation via repeated evolution, populations in similar habitats often differ substantially in phenotype and genotype (5–10). While easily dismissed as non-adaptive noise, such departures from convergence among populations from similar habitats may themselves reflect adaptation – not to shared selection pressures, but to unique local conditions within a given habitat type. Because such site-specific adaptation cannot be inferred from repeated evolutionary divergence, the same question arises for populations from similar habitats as for those compared across contrasting habitats: do their differences reflect local adaptation or not?

There are two broad explanations for why populations in similar habitats evolve differences (Figure 1A). First, differences may arise from non-adaptive processes such as drift (11–13), from alternative fitness-equivalent solutions to the same selection regime (14–16), or because populations vary in their ability to reach a shared fitness optimum owing to limited genetic variation, time, or homogenizing gene flow (17–21). In these cases, population variation within the same habitat type does not reflect adaptation to different local optima, but instead results from constraints or alternative adaptive responses when evolving toward a shared optimum (Figure 1B). Alternatively, ecological conditions may vary sufficiently among populations from the same nominal habitat to drive evolution toward different local fitness optima (3, 5, 6, 8, 22). In this case, population differences may reflect adaptation to site-specific selective conditions (Figure 1B). Although these explanations are not mutually exclusive, the extent to which differences among populations from seemingly similar habitats reflect adaptation to unique local conditions remains unresolved. More broadly, the relative contributions of adaptation to unique site-specific conditions and to shared habitat-level selection pressures for local fitness remain unclear. Resolving their relative contributions is critical: substantial site-specific adaptation would imply that even populations from similar habitats represent distinct evolutionary outcomes with independent conservation value. It would also suggest that much of what is often dismissed as non-adaptive noise is instead the outcome of natural selection shaped by fine-scale environmental variation.

**Figure 1.**
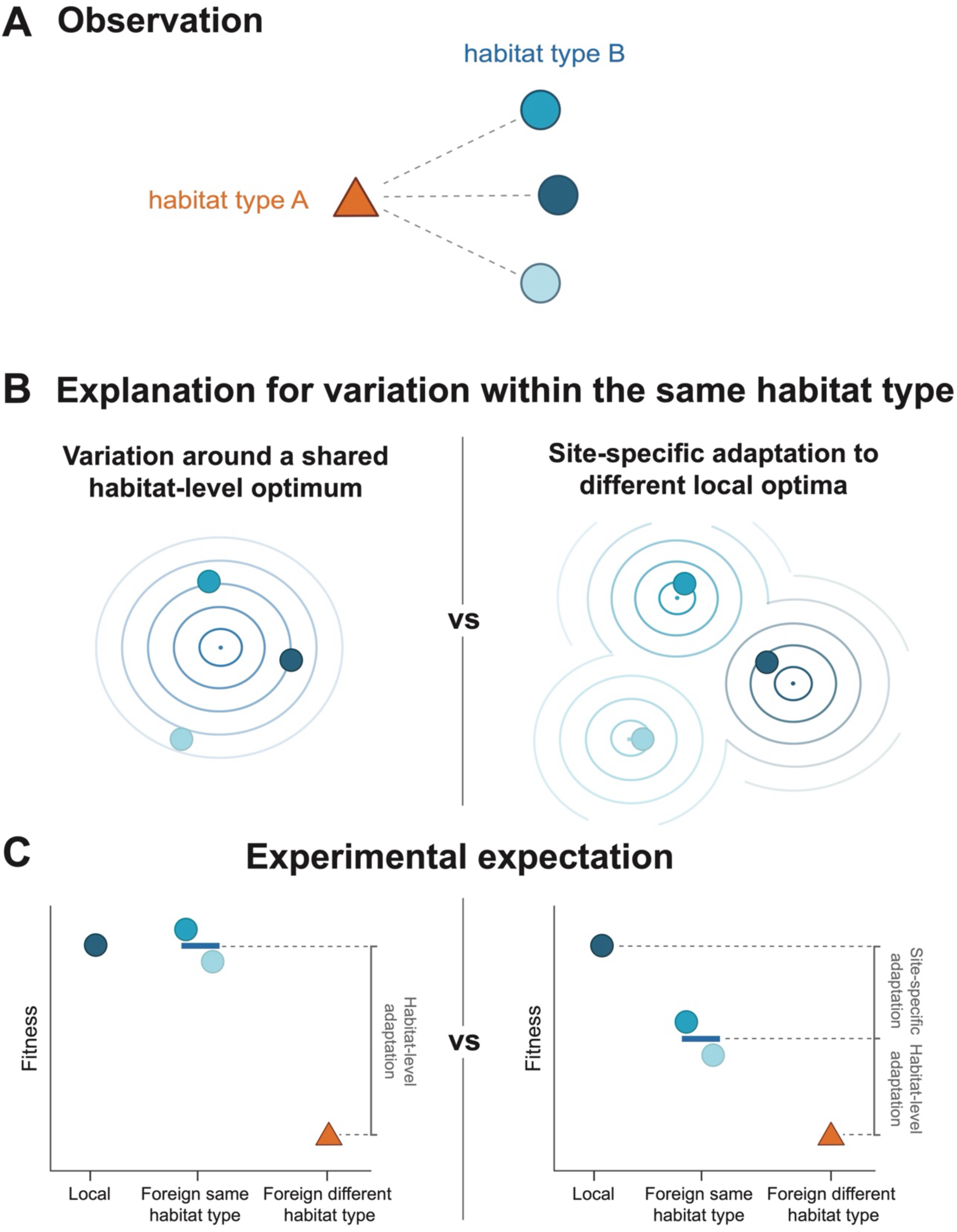
Conceptual framework for testing site-specific adaptation and evaluating its importance relative to shared habitat-level adaptation. (A) Repeated divergence between contrasting habitats (dotted lines) is commonly used to infer adaptation. However, populations from the same habitat type often differ from one another (circles with different blue tones), raising the question of whether such differences reflect site-specific adaptation within a given habitat type. (B) If all populations within a given habitat type experience similar selection pressures, they are expected to evolve toward a shared habitat-level fitness optimum, and differences among them could arise from non-adaptive processes, constraints, or alternative responses to the same selection regime (left). Alternatively, ecological conditions may differ sufficiently among sites within a given habitat type such that populations evolve toward distinct local fitness optima through site-specific selection (right). (C) Predictions for distinguishing these explanations using a transplant experiment involving multiple populations transplanted both within and across habitat types. At a focal site (illustrated for the dark blue population), the fitness of the local population is compared with that of foreign populations from the same habitat type and from a contrasting habitat type. If all populations from the same habitat type are adapted to a shared habitat-level optimum, the mean fitness of foreign populations from that habitat type (blue bar) is expected to match that of the local population, with no consistent local advantage (left). If populations are adapted to unique site-specific conditions, the local population is expected to exhibit higher fitness than the average of foreign populations from the same habitat type despite sharing a similar habitat-level selective regime (right). Including populations from a contrasting habitat type allows the fitness advantage of the local population over foreign populations to be partitioned into components attributable to shared habitat-level and unique site-specific adaptation.

How could we test whether populations from similar habitats are adapted to unique local conditions? One approach is to quantify ecological variation beyond broad habitat categories and determine how much this variation explains non-repeated population differences (e.g., 5, 8, 23–25). While informative, such tests remain inherently correlative and results hinge on the traits and environmental factors measured, and on assumptions about their relevance to selection. Thus, even when population differences covary with ecological variation, whether and how they translate into fitness in nature remains unknown. A more direct test of adaptation comes from transplant experiments that compare the performance of individuals from different populations under natural conditions (1, 26). By measuring whole-organism fitness directly in the wild, such experiments bypass assumptions about the specific traits or selective agents involved. Local adaptation is demonstrated when local individuals outperform foreign ones, indicating that natural selection favors the local type in its native habitat and drives divergence between populations (27–29). However, transplant experiments typically rely on pairwise reciprocal exchange between populations from contrasting habitats (1, 29, 30). Such pairwise comparisons can only quantify the overall strength of local adaptation between populations, without distinguishing how much reflects adaptation to the focal habitat contrast and how much reflects adaptation to unique conditions at individual sites. Consequently, they cannot determine whether populations from similar habitats are adaptively unique, nor can they quantify the relative contributions of site-specific versus habitat-level adaptation to overall fitness.

These limitations could be overcome by extending the classical reciprocal transplant design to include transplants of multiple populations not only across different habitats, but also among sites within the same habitat type. At a given site, this would allow comparing the fitness of the local population with that of foreign (allopatric) populations from both the same and different habitat types. These comparisons generate explicit, testable predictions (Figure 1C). If populations within a given habitat type are not adapted to distinct local optima, foreign populations from that same habitat type are expected to perform, on average, similarly to the local population. By contrast, if populations are adaptively tuned to site-specific ecological conditions, local individuals should exhibit higher mean fitness than foreign individuals, even when those originate from the same nominal habitat type. Importantly, including populations from a contrasting habitat does more than provide a positive experimental control. It allows site-specific adaptation to be quantified relative to shared adaptation, thereby separating responses to unique local selection pressures from those to common habitat-level selection pressures. Specifically, fitness differences between local and foreign populations from the same habitat type capture adaptation to unique local conditions, whereas fitness differences between foreign populations from the same versus a contrasting habitat type capture adaptation to selection pressures shared among populations from the same habitat type (Figure 1C). Together, these contrasts allow fitness variation among populations to be decomposed into effects arising from site-specific adaptation and from repeated responses to shared selection pressures.

Here, we applied this experimental framework in threespine stickleback fish *(Gasterosteus aculeatus)*. Following the retreat of glaciers from the last ice age about 12,000 years ago, marine stickleback repeatedly colonized countless lakes and streams across the Northern Hemisphere (31, 32). Extensive work has documented repeated divergence between lake and stream populations, consistent with divergent adaptation and thus repeated adaptation to shared selection pressures within habitat types. Yet populations from the same habitat type often differ substantially in phenotype and genotype (e.g., 6, 8, 10, 33–35). Whether such non-parallel variation reflects adaptation to unique local conditions, however, remains experimentally untested. In this study, we specifically focus on threespine stickleback from the Lake Constance watershed in western Europe, where a well-mixed lake population with a pelagic (open-water) lifestyle and morphology contrasts with multiple inlet populations adapted to benthic stream habitats. As elsewhere, previous work in this system has mainly focused on patterns of repeated lake–stream divergence (36–40), yet independent stream populations themselves show substantial variation (Figure 2A). This variation motivates an experimental test of whether these populations are adaptively unique and, if so, how much unique site-specific adaptation contributes to overall local adaptation compared with adaptation to shared habitat-level selection pressures.

**Figure 2.**
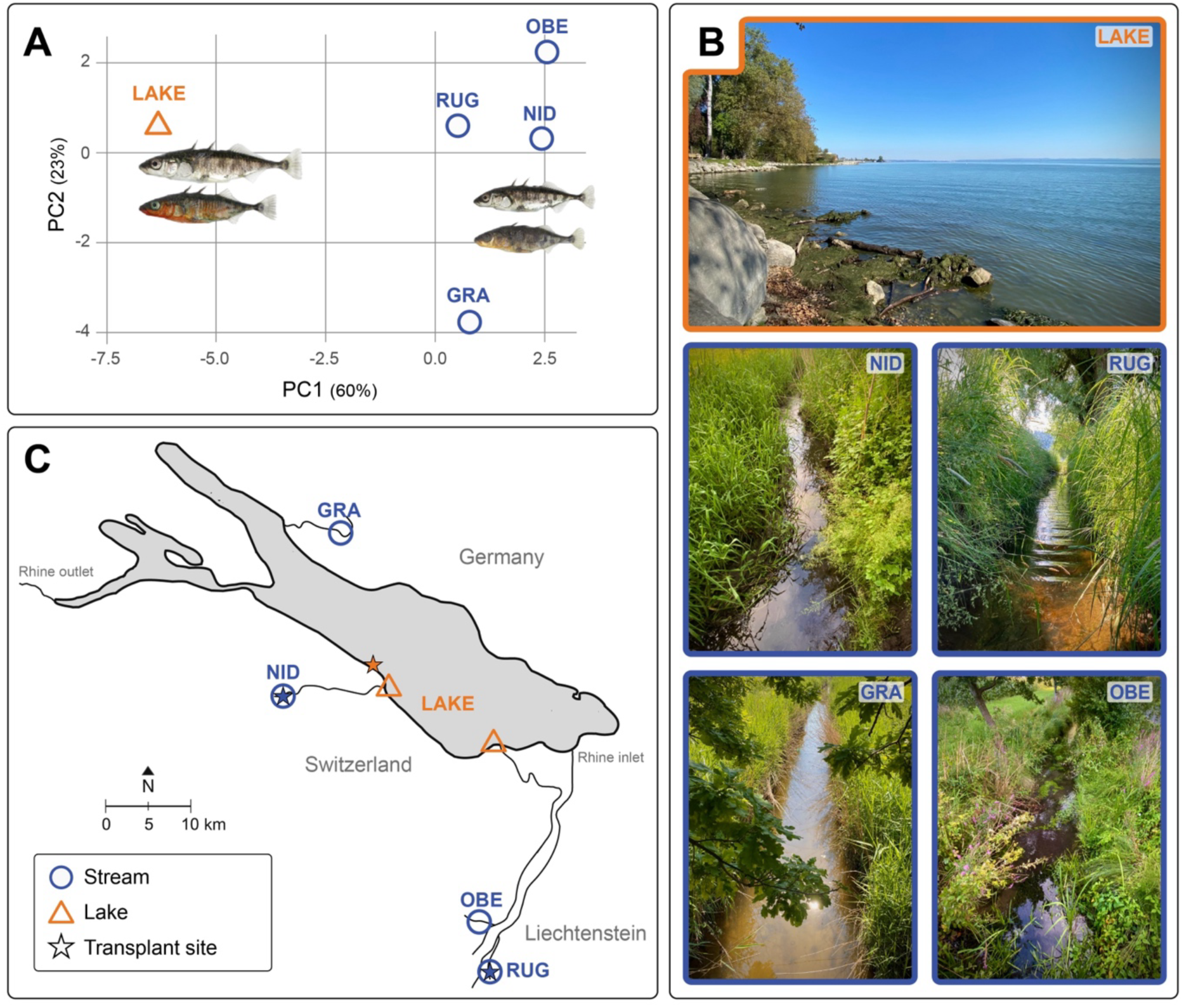
Study system and sites. **(A)** Stickleback in the Lake Constance watershed vary not only between lake and stream habitats but also among streams, potentially reflecting site-specific adaptation within the same habitat type. Shapes represent population means along the first two principal components of 22 morphological traits. The lake population is shown as an orange triangle and stream populations as blue circles. Inset: representative wild-caught females (top) and males (bottom) from the lake population (left) and the NID stream population (right). (**B**) Natural habitats of the focal lake and stream populations. (**C**) Map of the Lake Constance watershed with the location of the four stream populations (blue circles), the two sampling locations of the panmictic lake population (orange triangles), and the three experimental transplant sites (stars). Photographs of the experimental sites and enclosures are shown in Supplementary Figure S1.

## Results

To test whether populations inhabiting ecologically similar habitats are adaptively unique, and to quantify the relative contributions of site-specific and shared components of local adaptation, we conducted a multi-population field transplant experiment using threespine stickleback from lake and stream populations within the Lake Constance watershed (Figure 2B). Specifically, we transplanted individuals from pure within-population crosses originating from the panmictic lake population and four independent stream populations into field enclosures at one lake site and two independent stream sites (Figure 2C, Supplementary Figure S1). Fish were reared for two generations under common-garden laboratory conditions before transplantation, allowing evolved differences in fitness to be quantified in the wild. Each field enclosure received a balanced mixture of five lake fish and eight stream fish (two from each stream population), including individuals from the local population at each experimental field site. In total, we stocked 715 individually marked fish into 55 enclosures across the three field sites. After approximately seven weeks of exposure to natural conditions, we measured fitness using survival and body condition and estimated local adaptation as the average relative fitness advantage of local over foreign fish (28, 29).

Overall survival varied across the three experimental sites, being highest at the lake site (88%) and lower at the two stream sites (83% at NID and 55% at RUG). Importantly, within experimental sites, survival by fish type was consistent with both divergent lake–stream adaptation and site-specific adaptation (Figure 3A, Supplementary Table S1). That is, lake fish showed higher survival than stream fish at the lake site, whereas lake fish showed lowest survival at both stream sites. Moreover, at both stream sites, local stream fish showed higher survival than foreign stream fish. Although survival is a key component of fitness, especially before sexual maturity, body condition provides complementary information on an individual’s energetic state and expected future survival and reproductive success (41, 42). Across all fish, body condition was highest at the lake site and lower at the stream sites, thus mirroring the overall pattern observed for survival and indicating stronger overall performance of fish in the lake enclosures (Figure 3B). Body condition differences among fish types were weaker than for survival but showed the same pattern: lake fish outperformed stream fish at the lake site, stream fish outperformed lake fish at stream sites, and local stream fish outperformed foreign stream fish at both stream sites (Figure 3B). Overall, these consistent patterns across different fitness measures and field sites support adaptive lake–stream divergence with repeated adaptation to shared stream selection pressures as well as site-specific adaptation to unique local conditions.

**Figure 3.**
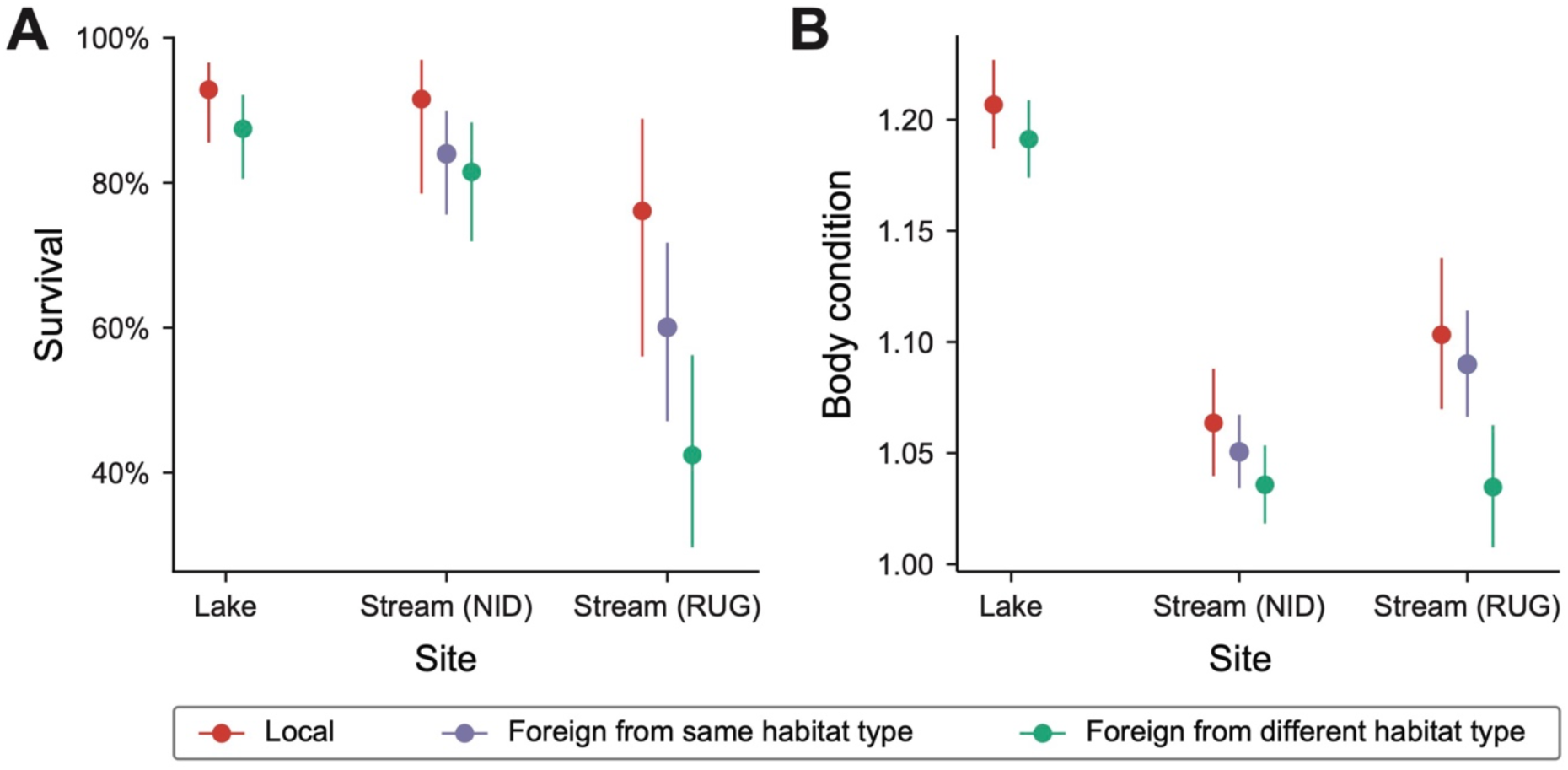
Fitness of different fish types across lake and stream transplant sites. **(A)** Survival and **(B)** body condition (Fulton’s index) of common-garden-raised threespine stickleback fish transplanted into one lake site and two independent stream sites. Fish were local or foreign relative to the transplant site, with foreign fish originating either from the same habitat type or from a different habitat type (lake vs. stream). At the lake site, local lake fish showed higher survival and body condition than foreign stream fish, whereas at both independent stream sites, local stream fish showed higher values than foreign lake fish. At both stream sites, local stream fish also showed higher survival and body condition than foreign stream fish. Points and error bars represent model-predicted means ± 95% confidence intervals.

Following the framework of classical reciprocal transplant experiments, we next used pairwise population contrasts to analyze fitness. For each contrast, both populations were tested at their local site of origin and at the foreign site of the other population. This allowed reciprocal comparisons both between ecologically divergent habitats (lake–stream) and between populations from the same habitat type (stream–stream). Across the two lake–stream contrasts (Lake–NID and Lake–RUG), local fish consistently outperformed foreign fish for both fitness components (Figure 4A; see Supplementary Figure S2 for individual contrasts). For survival, local–foreign differences were pronounced, with an average local fitness advantage of 16%, and were stronger at the stream sites. For body condition, local fish also showed consistent advantages, though of smaller magnitude, averaging 3% across sites. The reciprocal stream–stream comparison (NID–RUG) also revealed a clear local-over-foreign fitness advantage, with a 27% advantage for survival and a 2% advantage for body condition (Figure 4B). Together, these reciprocal contrasts indicate that local adaptation in this system arises both from divergence across the lake–stream divide and from site-specific adaptation within streams.

**Figure 4.**
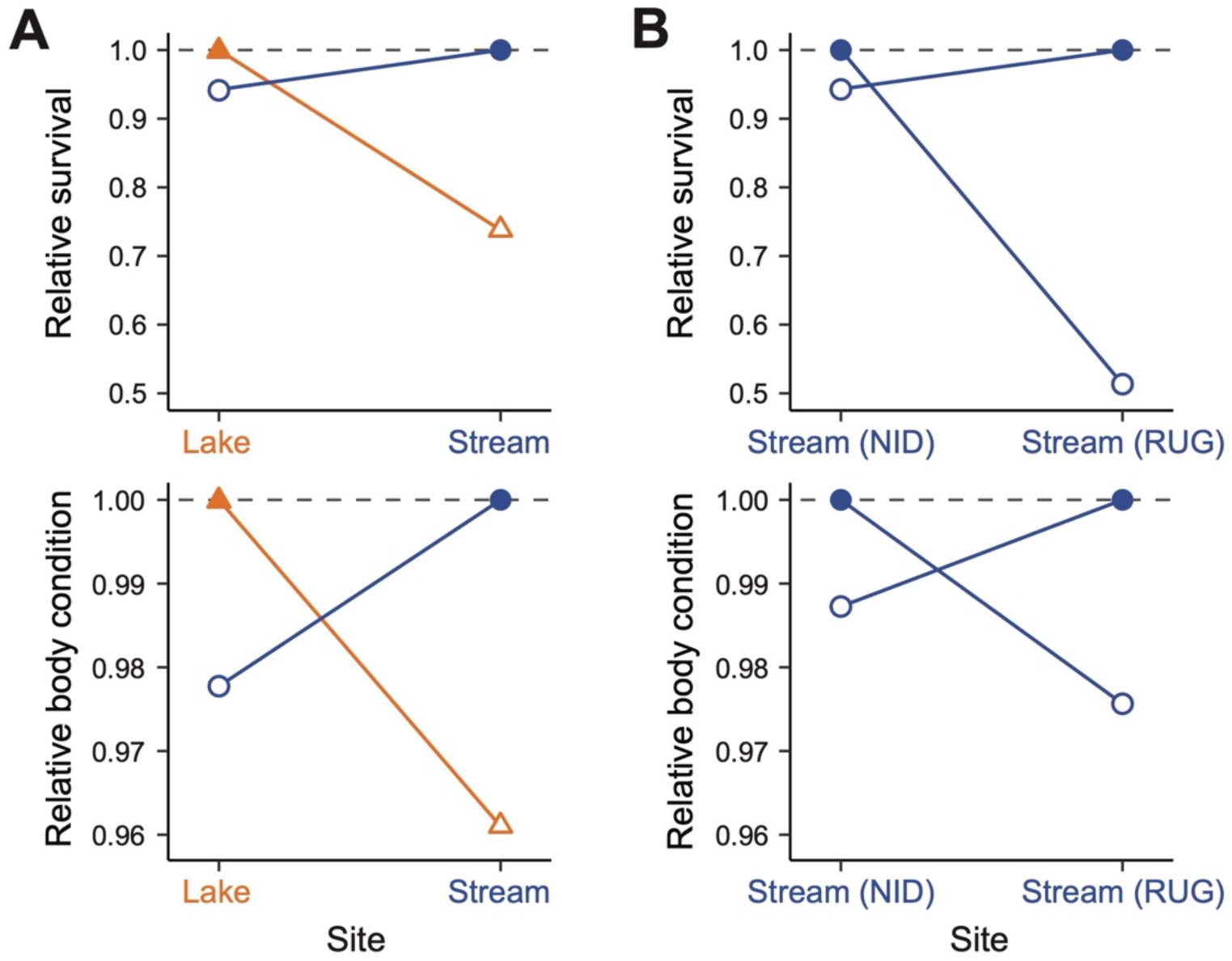
Reciprocal population contrasts reveal a consistent local-over-foreign fitness advantage. Pairwise reciprocal transplant comparisons showing relative fitness of local populations (filled symbols) and foreign populations (open symbols), standardized within each site such that the predicted fitness of the local population equals 1 (dashed horizontal line). Blue circles indicate stream populations and orange triangles indicate the lake population. (**A**) Mean reciprocal lake–stream contrasts, averaged across the Lake–NID and Lake–RUG comparisons, shown for survival (top) and body condition (bottom). (**B**) Reciprocal stream–stream contrast between the NID and RUG stream populations. Across fitness components and contrasts, foreign populations consistently show lower relative fitness than the local population at each site (values < 1), resulting in crossing reaction norms across sites. Plots are based on mixed-model predictions from data subsets including only the two populations involved in each reciprocal comparison.

To partition fitness variation into shared habitat-level and unique site-specific components, we focused on the two stream sites. The fitness advantage of foreign stream fish over foreign lake fish reflects adaptation to selection pressures shared across streams, whereas the fitness advantage of local stream fish over foreign stream fish reflects adaptation to site-specific conditions. To better capture and evaluate overall fitness, we combined model-predicted survival and body condition for each fish type into a composite fitness metric. Across both stream sites, composite fitness was highest for local stream fish (0.964), intermediate for foreign stream fish (0.788), and lowest for foreign lake fish (0.688). Observed values for local stream and foreign lake fish fell well outside the distributions expected under random assignment of fish types (both *P* < 0.002) (Figure 5). Local stream fish had 40% higher fitness than foreign lake fish (*P* < 0.001) and 24% higher fitness than foreign stream fish (*P* = 0.007), whereas foreign stream fish had 13% higher fitness than foreign lake fish (*P* = 0.067). Together, these contrasts allowed “total local adaptation” in streams – defined as the fitness advantage of local stream fish over foreign lake fish – to be decomposed into shared habitat-level and unique site-specific components. Indeed, reciprocal lake-stream contrasts confirmed that lake fish are locally adapted to their native habitat (Figures 3 & 4). Consequently, their reduced fitness in streams provides a valid baseline against which stream-specific fitness gains could be evaluated. Of the total fitness advantage of local stream fish over foreign lake fish, 33% reflected adaptation to selection pressures shared among streams (foreign stream vs. foreign lake), whereas 67% reflected site-specific adaptation to unique local conditions (local vs. foreign stream) (Figure 5). Thus, although adaptation to selection pressures shared among streams contributed substantially to local adaptation, most of the fitness advantage arose from unique site-specific adaptation.

**Figure 5.**
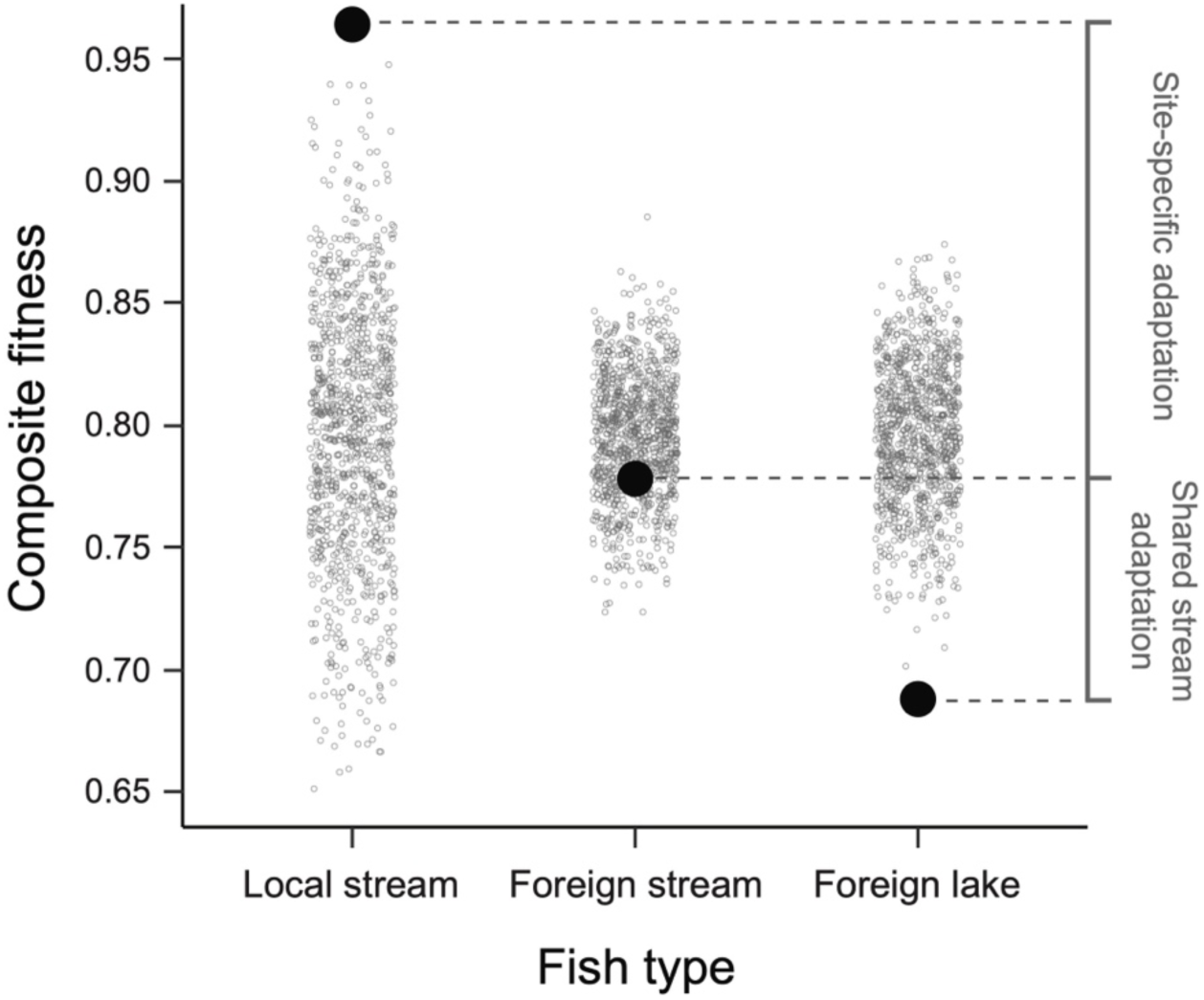
Quantifying the fitness effects of unique site-specific and shared habitat-level adaptation. Large filled circles show the observed composite fitness of local stream fish, foreign stream fish, and foreign lake fish across the two experimental stream sites. Composite fitness was calculated by integrating model-predicted survival and body condition. Small open circles show the permutation-based null distribution generated by randomizing fish-type labels within enclosures 1,000 times. Local stream fish lie above and foreign lake fish below the null distribution, indicating higher- and lower-than-expected composite fitness, respectively. Foreign stream fish show intermediate fitness, exceeding that of foreign lake fish but falling below that of local stream fish, and therefore overlap with the null distribution. This pattern indicates that foreign stream fish benefit from adaptation to selection pressures shared across stream habitats but lack adaptation to the unique site-specific conditions of the local population. On the right, the total fitness difference between local stream and foreign lake fish is partitioned into contributions from site-specific adaptation (local stream vs. foreign stream) and adaptation to selection pressures shared among streams (foreign stream vs. foreign lake).

## Discussion

Comparative studies typically infer local adaptation from repeated population divergence across clear ecological contrasts, yet replicate populations from similar habitats often differ in phenotype and genotype. While often dismissed as non-adaptive noise, such variation may instead reflect adaptation to unique ecological conditions among sites within the same nominal habitat. Ultimately, interpreting this variation and evaluating adaptation requires experimental tests of whole-organism fitness in nature.

We conducted such an experiment using lake and stream threespine stickleback from a single watershed, comparing the performance of local individuals with that of foreign individuals from the same and a contrasting habitat type. At both experimental sites in independent streams, we uncovered consistent evidence for site-specific adaptation: local stream fish outperformed foreign stream fish. Strikingly, this local fitness advantage was twice as large as the fitness advantage from adaptation to selection pressures shared across streams. At the same time, our study revealed local adaptation across the lake–stream divide: stream fish outperformed lake fish in streams, whereas lake fish outperformed stream fish in the lake. This reciprocal pattern confirmed that lake fish are locally adapted to their native habitat and thus provide a valid baseline within streams for interpreting the higher performance of foreign stream fish as evidence of general stream adaptation. Together, these results show that even among populations occupying the same nominal habitat, a substantial – and in our system dominant – portion of the local fitness advantage arises from adaptation to unique local selection pressures rather than to selection pressures shared at the habitat level.

These results have important implications for classical transplant studies, which test for local adaptation by reciprocally transplanting pairs of populations across contrasting habitats (1, 28–30). Because such experiments compare only two populations at a time, they estimate local adaptation by integrating all sources of fitness divergence between those populations. As a result, they cannot distinguish adaptation associated with the focal habitat contrast from adaptation to unique local conditions at individual sites. Our multi-population transplant experiment, incorporating comparisons both across and within habitat types, overcomes this limitation by allowing us to partition “total local adaptation” into these components, suggesting that fitness differences measured in classical transplant experiments may often reflect site-specific adaptation as much as – or even more than – adaptation to the focal habitat contrast itself.

Our findings also have important implications for comparative studies, which typically infer adaptation from repeated divergence across discrete habitats or ecotypes (1, 3, 4, 10). Our experimental framework incorporates the same habitat-based classification of populations that underlies many such studies. Indeed, the lake–stream classification has been widely used to study adaptation not only in sticklebacks (e.g., 6, 8, 33–40, 43–46), but also across many other fish and non-fish taxa (47–59). More broadly, grouping populations into discrete habitats or ecotypes is common across a wide range of ecological contrasts, including benthic versus limnetic forms, low-versus high-altitude environments, coastal versus inland habitats, environments with low versus high predation or competition, and alternative host associations (7, 60). Although such habitat-based classifications provide a practical framework for identifying adaptive traits and genomic regions in comparative studies, our results indicate that these classifications are often too coarse to capture a substantial component of adaptive divergence among populations.

Together, these insights help explain previously documented patterns in lake–stream stickleback and place them into a broader evolutionary context. Although extensive comparative phenotypic, ecological, and genomic work supports repeated divergence across the lake–stream divide, stickleback populations from the same habitat type often differ substantially in phenotype and genotype (e.g., 6, 8, 10, 33–35). At the same time, direct experimental evidence for divergent local adaptation between lake and stream habitats in nature has remained limited and circumstantial (61). Previous reciprocal transplant studies often report weak or asymmetric patterns of local adaptation (62–65). While these outcomes may reflect true biological variation, they can also arise from common experimental limitations such as the use of wild-caught fish, relatively short experimental durations, or limited power resulting from small or asymmetric sample sizes or from sparse experimental replication. For example, the only previous transplant study using fish from standardized crosses found higher performance of stream fish in streams relative to lake fish (66). However, the study lacked the reciprocal test in the lake and the inclusion of foreign populations from the same habitat type, limiting its ability to demonstrate divergent adaptation and to distinguish shared habitat-level from unique site-specific adaptation. Indeed, asymmetric performance observed in a single habitat is insufficient to demonstrate divergent local adaptation because one population may perform well across multiple habitats rather than being specifically adapted to its native habitat (28, 67, 68). Demonstrating divergent adaptation therefore requires reciprocal designs. Our reciprocal lake–stream transplant experiment, conducted with common-garden fish, provides direct evidence for heritable lake–stream adaptation. At the same time, the inclusion of multiple populations from the same habitat type revealed substantial adaptive variation beyond that associated with the lake–stream contrast, revealing how much adaptation is overlooked when inference is restricted to repeatable divergence across this ecological axis.

Accordingly, a strong focus on repeatability – although powerful for detecting shared adaptive responses – can overlook important population-specific adaptation. A key feature of our approach is that it does not depend on assumptions about repeatability or on prior knowledge of specific traits or genetic variants underlying adaptation. While prior comparative insights may motivate applying our experimental framework, such knowledge is not a prerequisite. Indeed, comparative studies may preferentially focus on traits or genes that are already known or predicted to be involved in repeated adaptation, which can bias inference toward higher apparent repeatability. Measured repeatability may thus overestimate the extent to which local adaptation is shared among populations. Moreover, repeatability-based inferences of local adaptation can be misleading when populations evolve different phenotypic solutions to the same adaptive problem, or when distinct phenotypic or genetic architectures yield similar functional performance (2). By classifying multiple populations based on ecological context and testing whole-organism fitness in nature, our experimental approach distinguishes shared habitat-level and unique site-specific adaptation without requiring prior knowledge of the traits, genes, or selective agents involved, or the degree of repeatability itself. At the same time, detecting substantial site-specific adaptation can help guide future efforts to identify the ecological factors underlying local fitness differences.

If local adaptation is commonly dominated by unique site-specific responses, non-parallel phenotypic and genomic divergence should be the expectation, even in systems where the focus lies on repeated evolution. Thus, the finding that adaptation to shared habitat selection explained only a minor part of overall local adaptation may help explain why genome scans and phenotypic analyses often reveal substantial non-parallelism among populations from seemingly similar environments. Rather than being predominantly non-adaptive, non-parallelism may commonly arise from adaptation to site-specific ecological conditions. Viewing non-parallel divergence as often reflecting adaptation to site-specific ecological conditions also bears directly on ecological speciation, where reproductive isolation evolves as a by-product of divergent adaptation (1, 69, 70). Although variation in divergence along major ecological axes has been associated with the strength of reproductive isolation (71–73), the sources of this variation have remained unclear. Our results suggest that deviations from replicated divergence may often be ecologically driven, and that variation in reproductive isolation among populations may therefore be more closely tied to fine-scale local adaptation than previously recognized.

Finally, our findings also speak to the predictability of evolution and how it depends on ecological scale. Repeated divergence across coarse ecological contrasts reflects responses to shared selection pressures, whereas site-specific adaptation reflects responses to unique local ecological contexts. Both of these components of adaptation arise deterministically from natural selection and are therefore, in principle, predictable (13, 74–77). However, the predictability of evolution depends on the ecological resolution at which environments are defined and measured. At coarse scales, evolutionary change may often appear repeatable, whereas at finer scales, natural selection can generate a mosaic of distinct adaptive responses. Even populations from ostensibly similar habitats may therefore represent predictable yet distinct evolutionary trajectories with independent evolutionary and conservation significance. Viewed this way, evolutionary predictability lies not only in repeated outcomes, but also in the fine-tuned responses of populations to their unique local environments.

## Methods

### Study system and generation of experimental fish

This study focuses on threespine stickleback fish from two habitat types within the Lake Constance watershed in western Europe: the lake population inhabiting Lake Constance proper and four populations from separate tributary streams (NID, RUG, GRA, and OBE). Lake Constance is a large oligotrophic peri-alpine lake (surface area of the main basin: 536 km²; average depth: ∼100 m) characterized by extensive open-water habitat. In contrast, the four tributaries share typical characteristics of stickleback stream habitats both within the Lake Constance watershed and elsewhere: they are shallow channels only a few meters wide, with slow to near-stagnant flow and abundant marginal vegetation, often accompanied by patches of in-stream vegetation near the banks. Although natural stream systems inevitably differ in some respects, the focal streams share broadly similar habitat characteristics and contrast markedly with the lake habitat (Figure 2B).

Whereas Lake Constance stickleback form a single genetically homogeneous (panmictic) lake population (39), the four streams drain independently into the lake, are not directly connected to one another, and are separated by several kilometers of land (Figure 2C). Consequently, stickleback inhabiting these tributaries represent independently evolving populations. Although the precise colonization history of stickleback within the watershed remains uncertain, divergence of these populations must be relatively recent and cannot predate deglaciation of the Lake Constance basin following retreat of the Rhine Glacier approximately 12,000 years ago (78). Further details on study populations and sites are provided in Supplementary Table S2.

To characterize phenotypic population variation motivating the subsequent field transplant experiment, we sampled a total of 417 adult stickleback, including 135 individuals from the panmictic lake population (collected at two locations, ALT and ROM) and 66–82 individuals from each of the four stream populations. Fish were captured using unbaited minnow traps between May and July in 2019 and 2020, balancing the representation of both sexes within populations. Individuals were immediately euthanized using an overdose of MS-222, a fin clip was preserved in 95% ethanol, and bodies were fixed in 4% paraformaldehyde (Supplementary Analysis 1).

For the field transplant experiment, we generated second-generation common-garden fish. Transplant experiments often use wild-caught individuals for practical reasons, particularly when working with animals. However, testing for an evolutionary response to divergent natural selection – and thus the heritable component of local adaptation – requires individuals reared under common-garden conditions for at least one full generation (28). Such naïve individuals control for population differences arising from plastic or parental carryover effects (79, 80) and allow potentially confounding sources of variation, including age structure and sex ratio, to be standardized (29, 68). Accordingly, we used sticklebacks from pure within-population F₂ crosses reared under controlled laboratory conditions before release into the wild. To generate these fish, we repeated the field sampling procedure described above in 2021 to establish 39 within-population F₁ crosses between wild-caught fish: 13 from the lake population (7 ROM and 6 ALT) and 26 from the four stream populations (9 GRA, 6 NID, 7 OBE, 4 RUG). F₁ crosses were produced by in vitro fertilization in the field, with a unique male–female pair used for each cross.

Fertilized F₁ egg clutches were transported to the stickleback facility at the University of Bern (Switzerland) and raised separately in 100-L tanks under standardized conditions, including diet (a mix of benthic-like mysis and limnetic-like live artemia), water parameters (pH, conductivity, and temperature), and light regime, with food amounts per tank adjusted to the number of individuals. At approximately 1.5 years of age, F₁ adults were used to produce 36 purebred F₂ families. For each F₂ family, males and females from different F₁ crosses but originating from the same natural population were paired, thereby reducing potential inbreeding among closely related individuals. This resulted in 12 lake F₂ families (derived from grandparents sampled at the two lake locations: 5 from ALT and 7 from ROM) and 24 stream F₂ families, including 6 from GRA, 7 from NID, 7 from OBE, and 4 from RUG (Supplementary Table S3). Families were again raised separately by family in 100-L tanks under standardized conditions (see above).

A few weeks before the start of the field transplant experiment, the sex of all experimental F₂ individuals was determined from DNA extracted from a small regenerating caudal fin clip using a well-established PCR assay (81). Individuals were subsequently marked with a small subcutaneous elastomer tag placed adjacent to the first or second dorsal spine, positioned slightly to the right side for males and to the left side for females. Different elastomer colors were used to distinguish families according to the population origin of their grandparents. Because the Lake Constance population has been described as effectively panmictic with no detectable genetic population structure (39), individuals originating from the two lake sampling locations (ALT and ROM) were treated as originating from a single population and are in this paper collectively referred to as “lake fish”.

### Experimental sites and enclosures

To test for local adaptation and partition shared habitat-level versus unique site-specific components of adaptation, we conducted transplant experiments at three natural field sites: one lake site representing the habitat of the lake population within the Lake Constance watershed, and two stream sites in independent inlet tributaries (NID and RUG) corresponding to the origins of two focal stream populations with available experimental F₂ fish. Experimental enclosures were established within these lake and stream sites to capture natural conditions, using designs adapted from previous stickleback field experiments (e.g., 65, 66) (Supplementary Figure S1).

Lake enclosures were constructed from cuboid high-density polyethylene (HDPE) net cages (3.5 mm mesh size) mounted on aluminum tube frames. Each lake enclosure measured 2 × 1 m (2 m² bottom surface), stood 1.5 m tall, and was placed in littoral water approximately 1 m deep (±0.2 m, depending on position and water-level fluctuations). Lake enclosures were located in shallow near-shore habitat where lake sticklebacks breed and forage from approximately May through early August and were abundant during the experiment. Enclosures were fully closed, with a zipper above the waterline that allowed fish to be introduced and then sealed, while the mesh permitted free water exchange and entry of typical stickleback prey (zooplankton and macroinvertebrates). Enclosures were anchored to the substrate using rebars, with edges connected by cords to enhance stability, and were partially sheltered from waves by a nearby pre-existing concrete dam inside the water. We installed a total of 20 such lake enclosures.

At the two experimental stream sites, enclosures were adapted to the smaller and more heterogeneous stream habitat structure. Stream enclosures were built from 1-m-wide sheets of galvanized steel mesh (4-mm mesh size), secured with rebars driven into the streambed and banks. The mesh was pressed deeply into the soft substrate to seal the bottom and prevent fish from escaping. Each enclosure covered roughly 2 m² and extended from the bank toward the mid-channel while allowing natural water flow through and around the structure. The open bottoms provided access to the natural substrate and its benthic macroinvertebrate fauna, the primary food source for stream-dwelling sticklebacks (37, 82). Enclosures were tightly covered on top with sewn-on mesh netting, leaving only a small temporary opening for adding fish, which was sealed thereafter. Enclosures at each stream site were distributed along the same reaches from which fish had been collected, spaced several meters apart to maintain flow conditions and to capture local habitat variation across an approximately 200 m stream stretch. Water depth within the enclosures matched natural conditions (typically 0.2-0.6 m). The experiment included 21 enclosures at the NID and 14 at the RUG experimental stream site. Differences in enclosure number among experimental sites were due to habitat availability, logistical considerations, and permitting constraints; otherwise, the experimental design was fully balanced (see below).

Each field enclosure was stocked with 13 experimental F₂ fish: 8 stream fish (one male and one female originating from each of the four stream populations GRA, NID, OBE, and RUG) and 5 lake fish (2-3 from each of the two lake sampling locations, ALT and ROM, with 2 or 3 of each sex), in accordance with natural stickleback densities observed during repeated sampling of these populations over multiple years. Enclosures at each of the two experimental stream sites therefore received fish originating from the local stream population, three foreign stream populations, and the foreign lake population. At the experimental lake site, enclosures received fish from the panmictic local lake population and all four foreign stream populations. Siblings from the same F₂ family were never enclosed together and families were represented at equal proportions across the three experimental sites. Together, these criteria ensured a fully balanced design with respect to population, family, and sex. All remaining aspects of fish assignment to enclosures and sites were random. In total, this resulted in 715 experimental fish stocked across 55 enclosures.

### Executing the field experiment and measuring fitness

Experimental fish were approximately 8 months old when transplanted into the wild and showed no signs of sexual maturity at either the beginning or the end of the field experiment, such that our estimates of selection are restricted to a standardized pre-mature life stage. One day prior to release, all fish were individually measured for mass and standard length (0.75 ± 0.18 g and 39.8 ± 2.3 mm, respectively; mean ± SD), and a small regenerating caudal fin clip was collected and stored in ethanol. Fish were transported in enclosure-specific buckets and gradually acclimated to ambient water and outdoor conditions, after which they were released into the enclosures. Releases occurred on 20 June 2023 at the lake site and on 24 June 2023 at the two stream sites. Enclosures were subsequently left undisturbed but visually inspected every 1-2 weeks for structural integrity, damage, and proper closure. After approximately seven weeks, all surviving fish at the lake site were recovered on 8 August 2023 by removing the entire net enclosures, which were then carefully searched for fish. At the two stream sites, recovery was achieved by placing two to three minnow traps per enclosure over several days and checking them regularly by lifting the traps to ensure complete retrieval of experimental fish. Trapping at the stream sites was conducted between 10–13 August 2023, with a few additional individuals collected at the NID site a few days later. Following euthanasia with an overdose of MS-222, all surviving experimental fish were re-measured for body mass (once in the field and again later in the laboratory, with the mean value used for analysis) and standard length. We also collected another fin clip, stored it in pure ethanol, and preserved whole specimens in 10% formalin.

Because each experimental fish was uniquely identifiable by its elastomer tag and enclosure ID, fitness measurements and subsequent analyses could be performed at the individual level. We quantified survival and body condition as fitness components. Survival was coded as a binary variable (1 for alive recovered individuals; 0 otherwise). Body condition was quantified using Fulton’s body condition index (K), calculated as 100 × (mass in grams / standard length³ in millimetres), with higher values indicating greater mass relative to body length (42, 83). Body condition was calculated for each fish at the start of the experiment and again for all survivors at the end of the experiment.

### Qualitative inference of local adaptation

At the two experimental stream sites (RUG and NID), we compared fitness of local stream fish with foreign stream and lake fish. Although fish from two additional stream populations (GRA and OBE) were not transplanted back to their sites of origin, their inclusion increased replication of foreign stream fish at both stream sites, thereby strengthening inference about site-specific adaptation – defined here as the expectation that local stream fish outperform the mean performance of foreign stream fish across populations (28). In contrast, comparisons between foreign stream fish and foreign lake fish at stream sites quantified adaptation to selection pressures shared among streams, independent of site-specific effects. Inference about stream adaptation requires a reciprocal reference, which was possible at the lake site. The Lake Constance watershed contains a single lake with a well-mixed population; therefore, a single lake site was sufficient to test the reciprocal expectation that lake fish outperform stream fish in the lake while stream fish outperform lake fish in streams.

Each individual was classified into one of three “fish type” categories based on the relationship between its population of origin and the experimental site. “Local” referred to individuals whose population of genetic origin matched the experimental site (lake fish at the experimental lake site, NID fish at the experimental NID site, and RUG fish at the experimental RUG site). “Foreign from same habitat type” referred to individuals originating from a different population but the same habitat type as the experimental site, whereas “foreign from different habitat type” referred to individuals originating from the alternative habitat type. For example, at the experimental RUG site, non-RUG stream fish were classified as “foreign from same habitat type”, whereas lake-origin fish were classified as “foreign from different habitat type”. Fish at the experimental lake site were classified as either “local” (lake-origin) or “foreign from different habitat type” (stream-origin).

First, we compared within-site fitness differences among fish types across the three experimental sites (Figure 3). These initial models were used to qualitatively evaluate two predictions under local adaptation: (i) under divergent adaptation between lake and stream habitats, stream fish should outperform lake fish at stream sites, whereas lake fish should outperform stream fish at the lake site; and (ii) under site-specific adaptation within streams, fish from local stream populations should outperform fish from foreign stream populations (Figure 1). Survival (binary response) was analyzed using generalized linear mixed models, whereas log-transformed final body condition (continuous response) was analyzed using linear mixed models. Fish type, experimental site, and their interaction were included as fixed effects, allowing fitness differences among fish types to vary freely across sites in the models. Enclosure ID was added as a random effect to account for non-independence among individuals. For body condition, log-transformed pre-transplant body condition was included as a covariate to control for baseline fitness differences among individuals. Population of origin could not be included in either model and pre-transplant body condition could not be included in the survival model due to convergence issues and were therefore omitted. This omission was unlikely to affect qualitative conclusions because population identity is largely captured by the fish-type classification itself, and population representation was fully balanced across enclosures and sites. However, both variables were included in subsequent targeted models used to quantify the relative contributions of shared stream adaptation and unique site-specific adaptation across the two experimental stream sites (see below).

Second, we evaluated local adaptation using pairwise reciprocal population contrasts following the logic of classical reciprocal transplant experiments, in which local individuals are expected to outperform foreign individuals at their site of origin (28, 29). Unlike classical reciprocal transplants, our design enabled reciprocal comparisons not only between populations from contrasting lake–stream habitats (lake–NID and lake–RUG), but also between populations from the same habitat type (NID–RUG). For each of these three pairwise comparisons, we subsetted the data to include only individuals from the two focal populations and fitted (generalized) linear mixed models with population of origin, experimental site, and their interaction as fixed effects, enclosure ID as a random effect, and log-transformed pre-transplant body condition as a covariate. Survival models were fitted to all individuals, whereas body-condition models were fitted only to survivors using log-transformed final body condition as the response. Evidence for local adaptation was assessed by testing whether fitness differences between populations changed sign across sites, as expected under local adaptation. To facilitate comparison across sites, population pairs, and fitness components, we standardized predicted fitness values within each site by the predicted fitness of the local population (local = 1). This removed differences in absolute fitness among sites and expressed performance in relative terms, consistent with the logic of classical two-population reciprocal transplant designs. For lake–stream contrasts, standardized estimates from the two independent comparisons (lake–NID and lake–RUG) were subsequently averaged to obtain a single estimate of lake–stream local adaptation per fitness component for presentation in the main text. The results for individual lake–stream contrasts are shown separately in Supplementary Figure S2.

### Quantifying shared and site-specific components of adaptation

Having established overall patterns of local adaptation across both fitness components, we quantified the relative contributions of shared habitat-level and unique site-specific adaptation. Because the Lake Constance watershed contains only a single panmictic lake population (and thus no foreign lake populations were available for transplantation), this partitioning could only be performed at stream sites. At each stream site, “total local adaptation” was defined as the fitness advantage of local stream fish over foreign lake fish. This total was then partitioned using contrasts that hold either habitat type or foreignness constant. Specifically, the average fitness advantage of foreign stream fish over foreign lake fish quantifies adaptation to selection pressures shared among streams, whereas the fitness advantage of local stream fish over foreign stream fish quantifies adaptation to unique local conditions at a given stream. Consequently, if both components contribute to total local adaptation, foreign stream fish are expected to exhibit intermediate fitness – higher than foreign lake fish but lower than local stream fish.

To estimate shared habitat-level and unique site-specific adaptation, we calculated a composite fitness metric integrating survival and body condition, which more effectively captures overall performance than either measure alone (29, 84, 85). Composite fitness was calculated as the product of predicted survival and predicted post-transplant body condition for each fish type (86). Predicted values were obtained from mixed models fitted separately for survival and body condition, with fish type as the focal explanatory variable, experimental stream site and log-transformed pre-transplant body condition as covariates, and enclosure ID and population of origin included as random effects. Estimated marginal means for each fish type were then multiplied to obtain composite fitness estimates. To assess statistical significance of the observed fitness variation across the three fish types, we used a permutation approach (87). Fish-type labels were randomly reassigned within enclosures to preserve enclosure-level structure, and the survival and body-condition models were refitted to each randomized dataset. Composite fitness was then recalculated for each such random permutation. A total of 1,000 permutations were performed and *P*-values were calculated by comparing observed composite fitness values and differences among fish types (local stream vs. foreign stream; foreign stream vs. foreign lake) to the corresponding null distributions generated by the permutations.

All analyses were conducted in *R v. 4.4.3* (88). Mixed-effects models were fitted using the *lme4* package. Estimated marginal means and model-based predictions were obtained using the *emmeans* package. Data handling and post-processing were performed using *dplyr*, and figures were produced using *ggplot2* based on model-predicted values, and verified using alternative visualization with *visreg*.

## Acknowledgments

We gratefully acknowledge all those who contributed to the preparation of experimental fish, provided logistical support, and assisted with fieldwork: Ruth Bauer, Fabrice Bouquet, Noelia Gerber, Leander Hollinger, Jelena Jäggi, Zuyao Liu, Dave Lutgen, Juliana Rodríguez-Fuentes, Carlos Rodríguez-Ramírez, Lotti Rösti, André Rösti, Axel Schmetzke, and Anine Wyser. We thank Anna Kropf for fish phenotyping and Daniel Berner for providing stickleback photographs for Figure 2A. We also thank Werner Bosshard, Jürg Fingerle, and Samuel Schmied for facilitating access to the experimental stream sites, and especially Beat Hirt for providing access to the experimental lake site. We further acknowledge the local fisheries, water, and veterinary authorities for approving and supporting this work: Christoph Birrer, Matthias Bopp, Werner Brunhart, Elke Desliens, Roland Jehle, Michael Kugler, Rainer Kühnis, Dario Moser, Mario Nutt, Jacques Voland, Markus Zellweger, and Marcel Zottele. M.R. would further like to extend his heartfelt thanks to Petra, Ronja, and Juna for their support and help behind the scenes during fieldwork. Finally, we are grateful to José Cerca, Jeffrey Groh, and Yoel Stuart for their valuable comments on the manuscript. This work was supported by the University of Bern and the Burgergemeinde Bern (grant to M.R.).

## Author Contributions

M.R. conceived and led research; M.R. conducted fish sampling; M.R., V.S., and N.N. made crosses, conducted fish husbandry, and prepared fish for the field experiment; M.R., H.R., and H.S. conducted the field experiment; C.L.P. and M.R. provided funding and other resources; M.R. analyzed the data and wrote the paper, with C.L.P., H.R., and H.S. providing feedback on the manuscript.

## Animal Care and Ethics

All fish sampling, husbandry, and field experimentation were conducted in accordance with applicable ethical, veterinary, and fisheries regulations (permit numbers: VTHa-Nr. BE4/16, BE116/2022, 664/2023-18155, LI-8471, SG-26.04.2021, TG-20.04.2020, and TG-30.09.2022).

## Data, Materials, and Software Availability

The code and raw data underlying this study are available on Dryad (DOI: XXXXX).

