## Supplementary Text, Figures, and Tables for "Disentangling unique site-specific and shared habitat-level adaptation in a classic system of repeated evolution"

### Supplementary Analysis 1

To illustrate phenotypic variation both between and, in particular, within habitat types that motivated the present field experiment, we phenotyped wild-caught stickleback from the focal populations within the Lake Constance watershed. In total, 417 individuals were phenotyped, including 66-82 fish from each of four stream populations and 135 fish from the panmictic lake population sampled at two locations. Sex representation was balanced within each population. Prior to phenotyping, fish were stained using an alkaline phosphatase protocol to facilitate later investigation of the lateral line system (for a separate study) and were subsequently stored in 10% buffered formalin (1).

A total of 22 phenotypic "traits" (mostly linear distances between fixed anatomical landmarks) were measured, many of which are known to vary with habitat or ecological niche in stickleback (2, 3) (Supplementary Analysis 1 Figure 1). Sixteen linear traits were measured from standardized digital photographs. Fish were pinned straight with the left side facing upward and photographed under controlled lighting. The left pectoral fin was removed prior to imaging. A total of 27 landmarks were digitized using tpsDig, including two landmarks corresponding to a 30-mm scale bar photographed with each specimen. Landmark coordinates (TPS format) were processed in *R* using the *geomorph* package and converted to millimeter scale using the reference length on each photograph, from which 16 linear distances and standard length (SL) were calculated. An additional five linear traits (gape width, buccal cavity length, pelvic girdle length, pectoral fin length, and epaxial muscle width) were measured directly on specimens using calipers, and the number of left-side lateral bony plates was counted under a dissecting microscope.

Traits were size-corrected using pooled within-group regression (4, 5). For each trait, an ANCOVA was fitted with group (population) and size (log-transformed SL) predicting the log-transformed trait value. Residuals from the common within-group slope were added to the predicted group trait value at the mean body size across all individuals, yielding size-adjusted values. To visualize among-population variation, a principal component analysis (PCA) was conducted on population trait means (5). Individual size-corrected trait values were averaged within populations, and the resulting means were analyzed using PCA on their correlation matrix (*prcomp()* in *R*), with results visualized using the *factoextra* package.

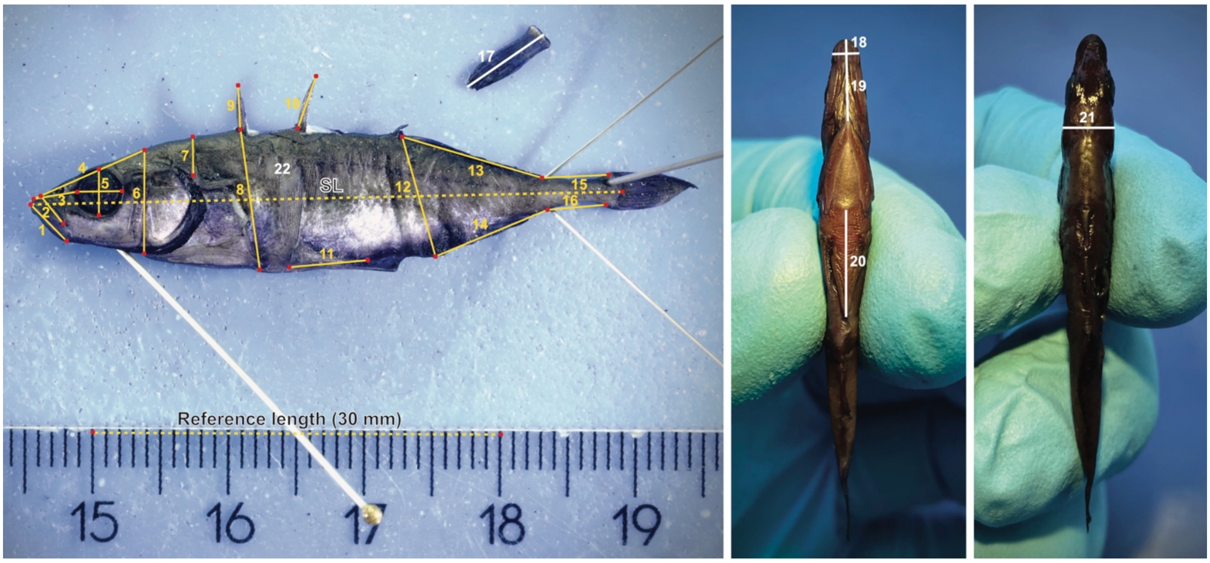


**Supplementary Analysis 1 Figure 1. Measured morphological traits.** Linear traits (indicated by yellow lines) were measured from digitized landmarks (red dots) placed on lateral images of fish. These traits included (yellow numbers): (1) jaw length, (2) mouth length, (3) snout length, (4) upper head length, (5) eye diameter (calculated as the mean of the two distances shown), (6) head depth, (7) epaxial muscle height, (8) anterior body depth, (9) first dorsal spine length, (10) second dorsal spine length, (11) pelvic spine length, (12) posterior body depth, (13) dorsal fin length, (14) anal fin length, (15) upper peduncle length, and (16) lower peduncle length. Four additional landmarks were used to determine standard length (SL) and a reference length (30 mm), which were used to scale all trait measurements. Additional traits were measured using handheld calipers (indicated by white lines): (17) pectoral fin length, (18) gape width, (19) buccal cavity length, (20) pelvic girdle length, and (21) epaxial muscle width. (22) The total number of left lateral body plates was counted under a microscope.

### Supplementary Figures

#### Supplementary Figure 1

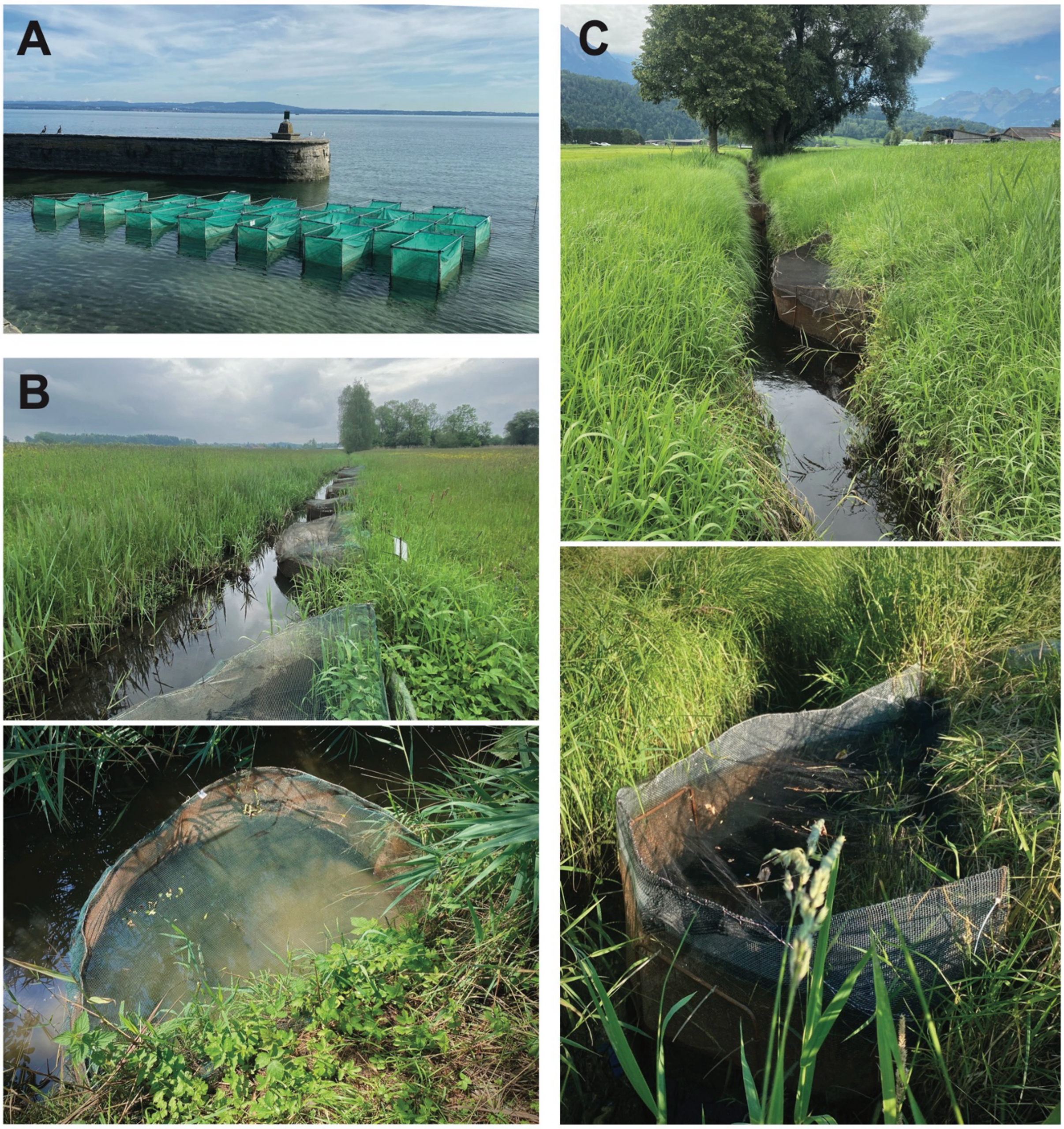


**Supplementary Figure 1. Experimental field sites and enclosures.** Enclosure designs in both lake and stream habitats were tailored to capture natural habitat variation while maintaining ambient environmental conditions. **(A)** Experimental lake site, consisting of 20 enclosures placed in ~1 m deep littoral water. **(B)** Snapshot of the experimental stream site in Niederaach (NID), with a close-up of one representative enclosure. **(C)** Snapshot of the experimental stream site in Ruggell (RUG), with a close-up of one representative enclosure.

**Supplementary Figure 2**

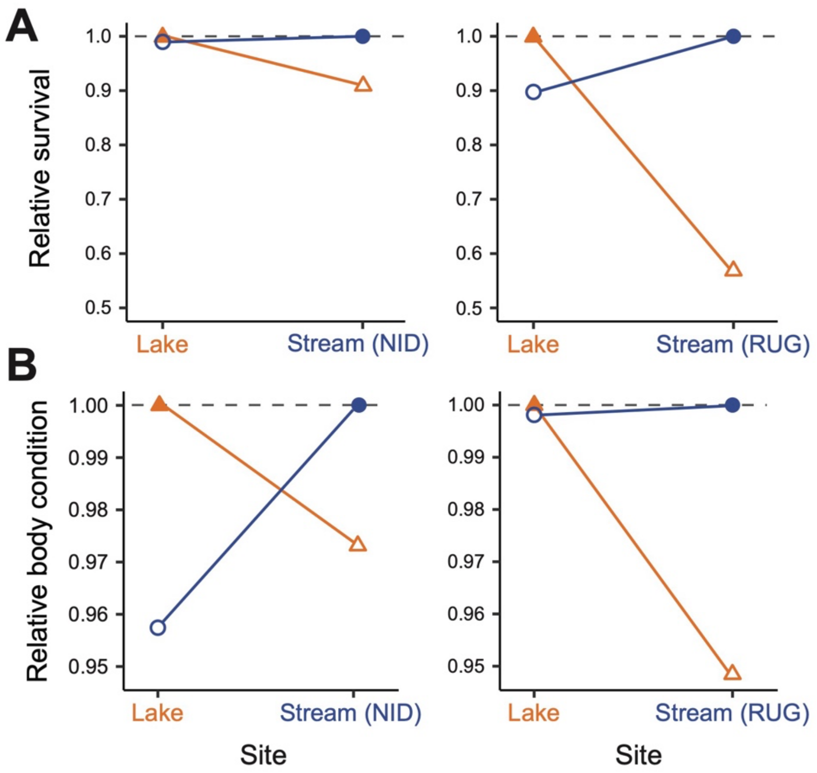


**Supplementary Figure 2. Reciprocal lake–stream contrasts reveal local-foreign fitness differences.** Pairwise reciprocal transplant comparisons showing predicted relative fitness of local and foreign populations, standardized within each site so the local population has relative fitness = 1 (dashed line). Filled symbols denote local populations and open symbols foreign populations; blue circles indicate stream populations and orange triangles the lake population. Comparisons are shown for Lake-NID and Lake-RUG population pairs for (**A**) survival and (**B**) body condition. Across fitness components and both lake-stream contrasts, foreign populations consistently have lower relative fitness than the local population (<1), producing crossing reaction norms between sites. Values are mixed-model predictions from datasets restricted to the two populations included in each reciprocal comparison.

### Supplementary Tables

**Supplementary Table S1. Survival frequencies.** Frequency of survivors for each fish type at each experimental site, with the number of survivors and total fish stocked shown in brackets.

| | | Experimental site | | |
|---|---|---|---|---|
| | | LAKE site | NID stream site | RUG stream site |
| Fish type | Local | **0.92** (92 / 100) | **0.90** (38 / 42) | **0.75** (21 / 28) |
| | Foreign same habitat type | - | **0.83** (104 / 126) | **0.60** (50 / 84) |
| | Foreign different habitat type | **0.86** (138 / 160) | **0.80** (84 / 105) | **0.43** (30 / 70) |

| **Supplementary Table S2. Details on study populations.** | | | | |
|---|---|---|---|---|
| **Population** | **Habitat** | **Details** | **latitude** | **longitude** |
| LAKE (ROM) | lake | Lake population sampled in Romanshorn (Switzerland) | 47.55607 | 9.38013 |
| LAKE (ALT) | lake | Lake population sampled in Altenrhein (Switzerland) | 47.48497 | 9.54331 |
| NID | stream | Salmsacher Aach sampled in Niederaach (Switzerland) | 47.54755 | 9.20839 |
| RUG | stream | Spiersbach sampled in Ruggell (Liechtenstein) | 47.25473 | 9.53824 |
| GRA | stream | Riedgraben sampled in Grasbeuren (Germany) | 47.71909 | 9.31563 |
| OBE | stream | Aubach sampled in Oberriet (Switzerland) | 47.31709 | 9.55585 |

**Supplementary Table S3. Overview of experimental fish.** $F_1$ families were produced from within-population crosses between wild-caught sticklebacks and reared under common-garden laboratory conditions. $F_2$ families were subsequently generated by crossing males and females originating from different $F_1$ families but from the same population, thereby producing purebred within-population $F_2$ families while minimizing potential inbreeding among closely related individuals. Pre-mature $F_2$ specimens were used in the field transplant experiment.

| **$F_2$ family ID** | **Habitat** [origin of grandparents] | **Population** [origin of grandparents] | **Date** [date $F_1$'s were crossed to obtain $F_2$ family] | **$F_1$ mother cross ID** | **$F_1$ father cross ID** | **No. experimental $F_2$ specimens** |
|---|---|---|---|---|---|---|
| LC.F2.20 | stream | GRA | 18.10.2022 | cross.38 | cross.33 | 17 |
| LC.F2.21 | stream | GRA | 18.10.2022 | cross.35 | cross.32 | 17 |
| LC.F2.22 | stream | GRA | 18.10.2022 | cross.39 | cross.40 | 22 |
| LC.F2.28 | stream | GRA | 31.10.2022 | cross.36 | cross.34 | 16 |
| LC.F2.36 | stream | GRA | 04.11.2022 | cross.36 | cross.34 | 21 |
| LC.F2.41 | stream | GRA | 05.11.2022 | cross.41 | cross.38 | 17 |
| LC.F2.01 | stream | NID | 17.10.2022 | cross.13 | cross.14 | 29 |
| LC.F2.02 | stream | NID | 17.10.2022 | cross.13 | cross.10 | 12 |
| LC.F2.03 | stream | NID | 17.10.2022 | cross.12 | cross.14 | 24 |
| LC.F2.04 | stream | NID | 17.10.2022 | cross.12 | cross.13 | 9 |
| LC.F2.05 | stream | NID | 17.10.2022 | cross.11 | cross.15 | 7 |
| LC.F2.06 | stream | NID | 17.10.2022 | cross.11 | cross.13 | 13 |
| LC.F2.14 | stream | NID | 17.10.2022 | cross.15 | cross.10 | 16 |
| LC.F2.23 | stream | OBE | 18.10.2022 | cross.22 | cross.01 | 16 |
| LC.F2.24 | stream | OBE | 18.10.2022 | cross.21 | cross.03 | 17 |
| LC.F2.25 | stream | OBE | 27.10.2022 | cross.03 | cross.05 | 12 |
| LC.F2.33 | stream | OBE | 03.11.2022 | cross.04 | cross.02 | 15 |
| LC.F2.38 | stream | OBE | 04.11.2022 | cross.05 | cross.21 | 16 |
| LC.F2.39 | stream | OBE | 04.11.2022 | cross.01 | cross.22 | 18 |
| LC.F2.40 | stream | OBE | 04.11.2022 | cross.22 | cross.04 | 16 |
| LC.F2.26 | stream | RUG | 27.10.2022 | cross.45 | cross.43 | 5 |
| LC.F2.31 | stream | RUG | 02.11.2022 | cross.43 | cross.42 | 21 |
| LC.F2.32 | stream | RUG | 02.11.2022 | cross.44 | cross.45 | 42 |
| LC.F2.42 | stream | RUG | 07.11.2022 | cross.42 | cross.44 | 42 |
| LC.F2.16 | lake | LAKE (ALT) | 18.10.2022 | cross.30 | cross.23 | 26 |
| LC.F2.29 | lake | LAKE (ALT) | 01.11.2022 | cross.31 | cross.25 | 34 |
| LC.F2.30 | lake | LAKE (ALT) | 02.11.2022 | cross.28 | cross.30 | 18 |
| LC.F2.35 | lake | LAKE (ALT) | 04.11.2022 | cross.24 | cross.31 | 32 |
| LC.F2.37 | lake | LAKE (ALT) | 04.11.2022 | cross.28 | cross.26 | 26 |
| LC.F2.07 | lake | LAKE (ROM) | 17.10.2022 | cross.17 | cross.16 | 32 |
| LC.F2.08 | lake | LAKE (ROM) | 17.10.2022 | cross.16 | cross.20 | 14 |
| LC.F2.09 | lake | LAKE (ROM) | 17.10.2022 | cross.19 | cross.17 | 17 |
| LC.F2.10 | lake | LAKE (ROM) | 17.10.2022 | cross.07 | cross.08 | 22 |
| LC.F2.11 | lake | LAKE (ROM) | 17.10.2022 | cross.08 | cross.19 | 24 |
| LC.F2.12 | lake | LAKE (ROM) | 17.10.2022 | cross.07 | cross.08 | 16 |
| LC.F2.13 | lake | LAKE (ROM) | 17.10.2022 | cross.19 | cross.16 | 14 |
| | | | | | **Total:** | **715** |
